# A dense brown trout (*Salmo trutta*) linkage map reveals recent chromosomal rearrangements in the *Salmo* genus and the impact of selection on linked neutral diversity

**DOI:** 10.1101/094763

**Authors:** Maeva Leitwein, Bruno Guinand, Juliette Pouzadoux, Erick Desmarais, Patrick Berrebi, Pierre-Alexandre Gagnaire

## Abstract

High-density linkage maps are valuable tools for conservation and eco-evolutionary issues. In salmonids, a complex rediploidization process consecutive to an ancient whole genome duplication event makes linkage maps of prime importance for investigating the evolutionary history of chromosome rearrangements. Here, we developed a high-density consensus linkage map for the brown trout (*Salmo trutta*), a socio-economically important species heavily impacted by human activities. A total of 3,977 ddRAD markers were mapped and ordered in 40 linkage groups using sex- and lineage-averaged recombination distances obtained from two family crosses. Performing map comparison between *S. trutta* and its sister species *Salmo salar* revealed extensive chromosomal rearrangements. Strikingly, all of the fusion and fission events that occurred after the *S*. *salar*/*S*. *trutta* speciation happened in the Atlantic salmon branch, whereas the brown trout remained closer to the ancestral chromosome structure. Using the strongly conserved synteny within chromosome arms, we aligned the brown trout linkage map to the Atlantic salmon genome sequence to estimate the local recombination rate in *S*. *trutta* at 3,721 loci. A significant positive correlation between recombination rate and within-population nucleotide diversity (π) was found, indicating that selection constrains variation at linked neutral sites in brown trout. This new high density linkage map provides a useful genomic resource for future aquaculture, conservation and eco-evolutionary studies in brown trout.

## INTRODUCTION

A renewed interest for linkage mapping studies has recently occurred thanks to the simplified procedures for generating large genotype datasets in non-model organisms (Slate *et al.* 2008; Stapley *et al.* 2010; Ekblom and Galindo 2011). High-density linkage maps have been particularly developed in teleost fishes, because of their high number of species, their economic importance for aquaculture (e.g., Sauvage *et al.* 2012; Zhao *et al.* 2013; Wang *et al.* 2015; Liu *et al.* 2016; Kumar and Kocour 2017), their phylogenetic position within vertebrates (Amores *et al.* 2011; Rondeau *et al.* 2014; Wang *et al.* 2015; Kuang *et al.* 2016), as well as their broad interests for eco-evolutionary and conservation/management issues (Everett *et al.* 2012; Gagnaire *et al.* 2013a; Kai *et al.* 2014).

The salmonid family, which has 11 genera and more than 60 species (Crête-Lafrenière *et al.* 2012), is a group of fish in which numerous initiatives have been developed to construct high-density linkage maps (Brieuc *et al.* 2014; Gonen *et al.* 2014; Limborg *et al.* 2015b; Larson *et al.* 2016; Tsai *et al.* 2016). Besides their applications for mapping traits of importance for aquaculture (Houston *et al.* 2012; Sauvage *et al.* 2012; Tsai *et al.* 2015), salmonid linkage maps have catalysed molecular ecology and genome evolution studies over the last recent years. Several studies have used linkage map information for addressing eco-evolutionary and related conservation biology issues at the species level or among hybridizing taxa (Lamaze *et al.* 2012; Bourret *et al.* 2013; Perrier *et al.* 2013; Gagnaire *et al.* 2013b; Limborg *et al.* 2014; McKinney *et al.* 2015). A second category of studies have focused on understanding the evolutionary consequences of the whole genome duplication event (salmonid-specific fourth vertebrate whole genome duplication; WGD-Ss4R) that occurred within the salmonid family around 60 Mya ago (Crête-Lafrenière *et al.* 2012), especially with regards to the partial rediploidization process which is still underway and not totally understood yet (Berthelot ei *al.* 2014; Brieuc *et al.* 2014; Kodama *et al.* 2014; Sutherland *et al.* 2016; Allendorf *et al.* 2015; Lien *et al.* 2016). A significantly improved understanding of this process has been achieved through increased efforts to map and compare chromosomal rearrangements among species (inversion, translocations, fissions and fusions), which helped to reconstruct the evolutionary history of salmonids genome architecture (Kodama *et al.* 2014; Sutherland *et al.* 2016). Furthermore, linkage mapping approaches have also improved the characterisation of the genomic regions that still exhibit tetrasomic inheritance patterns (Allendorf *et al.* 2015; Kodama *et al.* 2014; Lien *et al.* 2016; May and Delany 2015; Waples *et al.* 2016). The partial rediploidization of salmonid genomes is also known to challenge the development of genome-wide marker data sets. Indeed duplicated loci mostly located in tetrasomic regions (Allendorf *et al.* 2015) are usually underrepresent or even excluded in salmonid genomic studies (Hohenlohe *et al.* 2011; Hecht *et al.* 2012; Gagnaire *et al.* 2013b; Larson *et al.* 2014; Perrier *et al.* 2013; Leitwein *et al.* 2016)

The Eurasian brown trout (*Salmo trutta* L. 1758), is one of the most widespread freshwater species in the northern hemisphere, which presents high levels of phenotypic diversity linked to its complex evolutionary history (Elliott 1994; Bernatchez and Osinov 1995; Bernatchez, 2001). Natural brown trout populations have been intensely studied using molecular markers (Hansen *et al.* 2000; Sanz *et al.* 2006; Thaulow *et al.* 2014; Berrebi, 2015; Leitwein *et al.* 2016), but previous population genetics and phylogeography studies which usually relied on a limited number of markers did not integrate linkage map information (but see Hansen and Mensberg 2009; Hansen *et al.* 2010). The microsatellite linkage map available until now in brown trout (Gharbi *et al.* 2006) contains approximately 300 markers (288 microsatellites and 13 allozymes), which are distributed over 37 linkage groups (LGs) among the 40 LGs expected in this species (2*n*=80, Martinez *et al.* 1991; Philips and Ráb 2001). In order to follow the tendency towards an increasing number of markers in brown trout genetic studies (Leitwein *et al.* 2016), a higher density linkage map is needed to increase mapping resolution and provide positional information for genome scans and association studies. The development of a high-density linkage map in brown trout should find at least two direct applications.

Although the brown trout is the closest relative of the Atlantic salmon (*S*. *salar* L. 1758), the two species drastically differ in their number of chromosomes (*S*. *salar* 2*n* = 54-58, Brenna-Hansen *et al.* [2012]; *S*. *trutta* 2*n* = 80, Philips and Ráb [2001]), indicative of a high frequency of chromosome rearrangements. For understanding the chromosomal evolution in the *Salmo* genus, a comparison between the genome architecture of the Atlantic salmon (Lien *et al.* 2016) and the brown trout, using other salmonid linkage maps as outgroups, is needed. The availability of important genomic resources in salmonids coupled with recent methodological advances in linkage maps comparison (Sutherland *et al.* 2016) provide favorable conditions to get new insights into the recent history of chromosomal rearrangement in the *Salmo* genus.

Another important application of a high-density linkage map in brown trout relates to understanding the consequences of human-mediated introductions of foreign and domestic genotypes in natural populations. The brown trout has been domesticated for decades, and wild populations have been heavily impacted by the stocking of genetically domesticated hatchery strains sometimes originating from distinct evolutionary lineages than the recipient populations (Elliot 1994; Laikre 1999; Berrebi *et al.* 2000; Bohling *et al.* 2016). These introductions have often resulted in the introgression of foreign alleles within wild brown trout populations, a generally negatively perceived phenomenon in conservation and management (Largiadèr and Scholl 1996; Poteaux *et al.* 1998, 1999; Hansen *et al.* 2000; Almodóvar *et al.* 2006; Sanz *et al.* 2006; Hansen and Mensberg 2009). However, the real evolutionary consequences of hybridization may take multiple facets (e.g., outbreeding depression, heterosis, adaptive introgression, decreased adaptive variation), the individual effects of which remains largely unknown. In order to evaluate those potential fitness outcomes in brown trout, it would be necessary to combine the power of large SNP data sets (Leitwein *et al.* 2016) with a high-density linkage map and a detailed knowledge of chromosomal variation in recombination rate to perform comprehensive genome scans for introgression in natural populations.

The specific objectives of this study were: (i) to build a high-density linkage map in brown trout based on RAD markers developed by Leitwein *et al.* (2016) using reciprocal crosses of individuals belonging to two distinct evolutionary lineages (Atlantic and Mediterranean; Bernatchez, 2001); (ii) to characterize chromosome morphologies and centromere positions of metacentric chromosomes using the method recently developed in Limborg *et al.* (2015a); (iii) to perform map comparisons with the *S*. *salar* genome and an outgroup salmonid species to infer the timing of chromosome rearrangements during the recent evolutionary history of the *Salmo* genus, and (iv) to estimate the genome wide local variation in recombination rate and assess the consequence of this variation on the genomic landscape of nucleotide diversity in brown trout.

## MATERIALS AND METHODS

### Experimental crosses

Two F1 hybrid families, each comprising two parents and 150 offspring, were used for the linkage map analysis. Both were obtained by crossing one parent of Atlantic origin with one parent of Mediterranean origin *sensu* Bernatchez (2001). The F0 progenitors were either from a domestic Mediterranean strain lineage originally bred in the Babeau hatchery (described in Bohling *et al.* [2016]) and a domestic Atlantic strain lineage originating from the Cauterets hatchery and independently reared from the Mediterranean strain at the Babeau hatchery. The Cauterets strain is one of the common, internationally-distributed Atlantic strains (comATL category; Bohling *et al.* 2016). Crosses were performed in December 2015 at the Babeau hatchery as follows: FAMILY 1: one Atlantic female × one Mediterranean male, and FAMILY 2: one Atlantic male × one Mediterranean female. Progenies were sacrificed in February 2016 after full resorption of the yolk sac (fish size ~ 2cm).

### DNA extraction, library preparation and sequencing

Whole genomic DNA was extracted from caudal fin clips for the four parents of the two crosses and from tail clips for 150 offspring of each family using the commercial Thermo Scientific KingFisher Flex Cell and Tissue DNA kit. Individual DNA concentration was quantified using both NanoDrop ND-8000 spectrophotometer (Thermo Fisher Scientific) and Qubit 1.0 Fluorometer (Invitrogen, Thermo Fisher Scientific). Double digest RAD sequencing library preparation was performed following Leitwein *et al.* (2016). Briefly, DNA was digested with two restriction enzymes (*Eco*RI-HF and *Msp*I), the digested genomic DNA was then ligated to adapters and barcodes. Ligated samples with unique barcodes were pooled in equimolar proportions and cleanPCR beads (GC Biotech, The Netherlands) were used to select fragments size between 200 and 700 base pairs (bp). The size selected template was amplified by PCR and tagged with unique index specifically designed for Illumina sequencing (Peterson *et al.* 2012). For progenies, PCR products were pooled in equimolar ratios into 4 pools. Each pool was sequenced in a single Illumina Hiseq 2500 lane (i.e. 75 F1 progenies per lane), producing 125bp paired-end reads. The 4 parents were sequenced in a single Illumina lane in order to obtain a higher coverage depth.

### Genotyping pipeline

Bioinformatic analyses followed the same procedure as in Leitwein *et al.* (2016). Briefly, raw reads passing Illumina purity filter and FastQC quality were demultiplexed, cleaned and trimmed with `process_radtags.pl` implemented in Stacks v1.35 (Catchen *et al.* 2013). The Atlantic salmon genome (GenBank accession number: GCA_000233375.4_ICSASG_v2) was used to determine individual genotypes at RAD markers using a reference mapping approach. First, reads were aligned to *S*. *salar* genome with the BWA-mem program (Li and Durbin, 2010). Alignment result files were processed into pstacks to build loci and call haplotypes for each individuals (using m = 3 and α = 0.05). Individuals with a minimum mean coverage depth under 7X were removed, resulting in a data set of 147 offspring in FAMILY 1 and 149 offspring in FAMILY 2. Hereafter, a RAD locus is defined as a sequence of 120bp, and thus can contain more than one SNP defining different haplotypes. A reference catalog of RAD loci was built with `cstacks` from the four parents of the two crosses (FAMILY 1 and Family 2). Each individual offspring and each parent were then matched against the catalog with `sstacks`. Then, for each family independently, the module `genotypes` was executed to identify mappable makers that were genotyped in at least 130 offspring in each family, and with a minimum sequencing depth of 5 reads per allele. The correction option (-c) in the `genotypes` module was used for automated correction of false-negative heterozygote calls. The map type “cross CP” for heterogeneously heterozygous populations was selected to export haplotypic genotypes in the JoinMap 4.0 format (Van Ooijen, 2006). We only kept the loci shared between FAMILY 1 and FAMILY 2, for importation into JoinMap, leaving a total of 7680 mappable loci in both families.

### Linkage mapping

Both family data sets were imported and filtered into JoinMap 4.0 (Van Ooijen, 2006) to remove (i) loci presenting significant segregation distortion, (ii) loci in perfect linkage (i.e., similarity of loci = 1), and (iii) individuals with high rates of missing genotypes (>25%). After this first filtering step, 5,635 and 5,031 loci remained in FAMILY 1 and FAMILY 2 respectively. Only those that were shared by the two families were kept, resulting in a subset of 4,270 filtered informative loci. A random subset of 4,000 loci for both FAMILY 1 and FAMILY 2 were reloaded into JoinMap 4.0 (i.e., the maximum number of loci handled by the program). We then discarded 4 offspring in FAMILY 1 and 6 offspring in FAMILY 2 due to high rates of missing genotypes (above 20%). Markers were then grouped independently within each family using the “independence LOD” parameter in JoinMap 4.0 with a minimum LOD value of 10. Unassigned markers were secondly assigned using the strongest cross-linkage option with a LOD threshold of 5. Marker ordering was finally estimated within each group using the regression mapping algorithm and three rounds of ordering. The Kosambi’s mapping function was used to estimate genetic distances between markers in centimorgans (cM). Loci with undetermined linkage were then removed, and a consensus map was produced between families using the map integration function after identifying homologous LGs between FAMILY 1 and FAMILY 2.

### Estimation of centromere location and chromosome type

The method recently developed by Limborg *et al.* (2015a) was used to identify chromosome types (acrocentric, metacentric) and to determine the approximate centromere position of metacentric chromosomes. This method uses phased progeny’s genotypes to detect individual recombination events by comparison to parental haplotype combinations. For each LG, the cumulated number of recombination events between the first marker and increasingly distant markers was computed from both extremities, using each terminal marker as a reference starting point. We used the subset of markers that were found heterozygous only in the female parent to phase progeny’s haplotypes within each family. We only focused on female informative sites because the lower recombination rate in male salmonids makes them less adequate for inferring centromere locations (Limborg *et al.* 2015a). Lower recombination in male brown trout was reported by Gharbi *et al.* (2006). The phased genotypes data sets were used to estimate recombination frequency (RFm) for intervals between markers along each LG. Finally, we plotted the RFm calculated from each LG extremity to determine the chromosome type and the approximate location of the centromere for metacentric chromosomes.

### Map comparisons

The microsatellite linkage map established by Gharbi *et al.* (2006) was compared and anchored to our new high density linkage map using the MAPCOMP pipeline available at https://github.com/enormandeau/mapcomp/ (Sutherland *et al.* 2016). The microsatellite markers’ primer sequences retrieved from Gharbi *et al.* (2006), as well as the five microsatellites used in Leitwein *et al.* (2016) and the lactate dehydrogenase (LDH-C1) gene from McMeel *et al.* (2001) were aligned with `BWA_aln` (Li and Durbin 2010) to the Atlantic salmon reference genome. Only single hit markers with a quality score (MAPQ) above 10 were retained. The 3,977 RAD markers integrated in the high density *S*. *trutta* linkage map (see results) were also aligned to the *S*. *salar* reference genome with `BWA_mem`. Microsatellite and LDH markers were anchored onto the RAD linkage map by identifying the closest RAD locus located on the same contig or scaffold. Pairing with closely linked RAD markers enabled us to position markers from the previous generation map onto the newly developed linkage map.

An additional comparison was performed between the *S*. *salar* high density linkage map published by Tsai *et al.* (2016), and the *S*. *trutta* consensus linkage map developed in this study. We used the alignment of Atlantic salmon and brown trout markers to the Atlantic salmon reference genome obtained with `Bwa_mem` (Li and Durbin 2010) to identify loci originating from orthologous genomic regions, and then used these loci in MAPCOMP (Sutherland *et al.* 2016) to identify blocks of conserved synteny and orthologous chromosome arms between the two species’ maps. Conserved synteny blocks were visualized with the web-based VGSC (Vector Graph toolkit of genome Synteny and Collinearity: http://bio.nj_fu.edu.cn/vgsc-web/). The brook charr (*Salvelinus fontinalis*) linkage map published in Sutherland *et al.* (2016) was used as one outgroup to provide orientation for chromosome rearrangements during evolution of the *Salmo* lineage.

In order to estimate the genome-wide averaged net nucleotide divergence between *S*. *trutta* and *S*. *salar* (*d*_a_), we used the mean nucleotide diversity within *S*. *trutta* (π_w_, averaged between Atlantic and Mediterranean lineages) (Leitwein *et al.* 2016) and the absolute nucleotide divergence between *S*. *trutta* and *S*. *salar* (π_b_, also called *d*_xy_). The net divergence was estimated as *d*_a_ = π_b_ − π_w_, by supposing that the mean nucleotide diversity in *S*. *salar* (which is not available) was close to the mean nucleotide diversity estimated in *S*. *trutta*. To estimate π_b_, we used the 3,977 consensus sequences of the RAD loci integrated in our *S*. *trutta* linkage map, and aligned them with `BWA_mem` to the Atlantic salmon reference genome. Then, the total number of mismatches divided by the total number of mapped nucleotides was used to provide an estimate of π_b_.

### Estimation of genome-wide recombination rates

To estimate local variation in recombination rate across the genome, we compared our newly constructed genetic map of *S*. *trutta* to the physical map of *S*. *salar* with MAREYMAP (Rezvoy *et al.* 2007). In order to reconstruct a collinear reference genome for each brown trout linkage group, physical positions of brown trout RAD markers on the salmon genome were extracted from the `BWA_mem` alignment for each block of conserved synteny previously identified with MAPCOMP. The Loess method was used to estimate local recombination rates. It performs a local polynomial regression to fit the relationship between the physical and the genetic distances. The window size, defined as the percentage of the total number of markers to take into account for fitting the local polynomial curve, was set to 0.9 to account for local uncertainties due to the high rate of chromosomal rearrangements between species. A recombination rate equal to the weighted mean recombination rate of the two closest markers on the genetic map was assigned to the markers that were not used for fitting.

### Correlation between recombination rate and nucleotide diversity

In order to test for a correlation between local recombination rate and nucleotide diversity in brown trout, we used the estimate of nucleotide diversity (π) in the Atlantic lineage (based on 20 individuals). These estimates of nucleotide diversity were taken from Leitwein *et al.* (2016), in which brown trout populations and hatchery strains of different origins showed highly correlated genome-wide variation patterns in nucleotide diversity. Markers with a recombination rate value above 2 cM/Mb or π < 0.008 were considered as outliers and were therefore removed from the analysis. This resulted in a dataset containing 3,038 markers for which both recombination rate and nucleotide diversity information were available.

### Data availability

Table S1 contains the RAD markers IDs, the consensus sequences, the mapping position on the brown trout linkage map along with the position on the salmon genome and the estimation of the recombination rate. Figure S1 contains the RFm plots obtained for each female haploid data set. Table S2 contains the correspondence between the microsatellite and the new RAD linkage map. Figure S2 contains the MapComp figure comparing the Brown trout to the Atlantic salmon linkage map with markers paired through the Atlantic salmon genome.

## RESULTS

### Construction of a high-density linkage map in *Salmo trutta*

On average, 35M of filtered paired-end (PE) reads were obtained for each of the four parents (two families), and 5M PE reads for each of the 300 F1 (150 progenies per family) analyzed in this study. Therefore, the retained subset of 7,680 informative markers common to both families had a mean individual sequencing depth of 112X for the parents and 15X for the progenies, ensuring a good genotype calling quality in both families. Among these, a subset of 4,000 markers showing no segregation distortion in both families was randomly retained, to generate a consensus map between families 1 and 2. The resulting integrated map provides a sex- and lineage-averaged map, which is an important feature considering the reported differences in recombination rates between male and female in salmonids (Allendorf *et al.* 2015; Tsai *et al.* 2015), and the existence of divergent geographic lineages in brown trout (Bernatchez 2001).

A total of 3,977 markers were assigned to 40 *S*. *trutta* (*Str*) LGs on the integrated map (Figure 1), corresponding to the expected number of chromosomes in brown trout (Phillips and Ráb, 2001). The total map length was 1,403 cM, and individual LGs ranged from 14.6 cM (*St*r3; 47 markers) to 84.9 cM (*Str*37; 63 markers). The number of markers per LG ranged from 23 for *Str*40 (15.2 cM) to 252 for *Str*1 (46.3 cM). Detailed information for each LG including the density of markers per cM is reported in Table 1. Additional details concerning RAD markers IDs, consensus sequences and mapping position on the salmon genome are reported in Table S1.

**Fig. 1.**
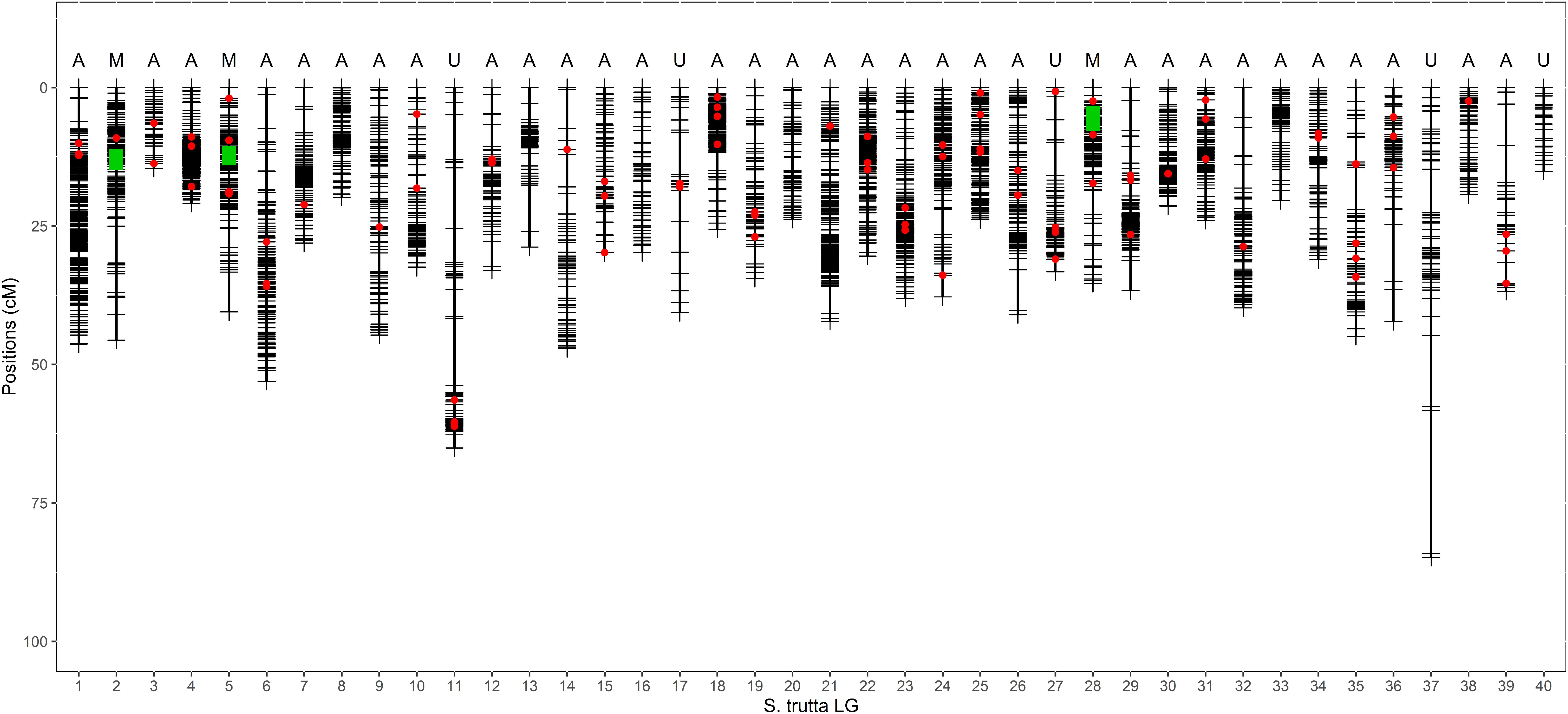
The brown trout (*Salmo trutta*) high density linkage map. Horizontal black lines within each of the 40 LGs represent mapped RAD markers (*n* = 3,977). The most probable chromosome type determined with the RFm method (Limborg et al. 2015a) is indicated above each LG (A: acrocentric, M: metacentric, U: undetermined). Green boxes indicate the approximate position of the centromere for metacentric LGs. Red dots represent the position of 89 additional markers that were transferred from the previous generation linkage map (see Table S2 for details).

### Location of centromeres and chromosome types

The estimates of recombination frequencies obtained using the phased genotypes of female progeny produced similar results for both FAMILY 1 and FAMILY 2. Individual linkage group RFm plots obtained for each female haploid data set are provided in Figure S1. Two examples of RFm plots obtained in FAMILY 1 are reported in Figure 2 for *Str*22 and *Str*28. Following Limborg *et al.* (2015a), the pattern observed for *Str*22 typically reflects an acrocentric chromosome type (paired straight lines; Figure 2a), whereas *Str*28 illustrates a metacentric pattern (mirrored hockey stick shapes; Figure 2b) with the centromere approximately located around 70cM. The total number of probable acrocentric chromosomes (Figure 1; type A, *n* = 32) largely exceeded the number of metacentric chromosomes (Figure 1; type M, *n* = 3). The approximate position of the centromere was determined for each of the three metacentric LGs (Figure 1; green boxes). The chromosome type remained undetermined for five LGs (Figure 1; type U) due to a low resolution of the RFm plots (Figure S1).

**Fig. 2.**
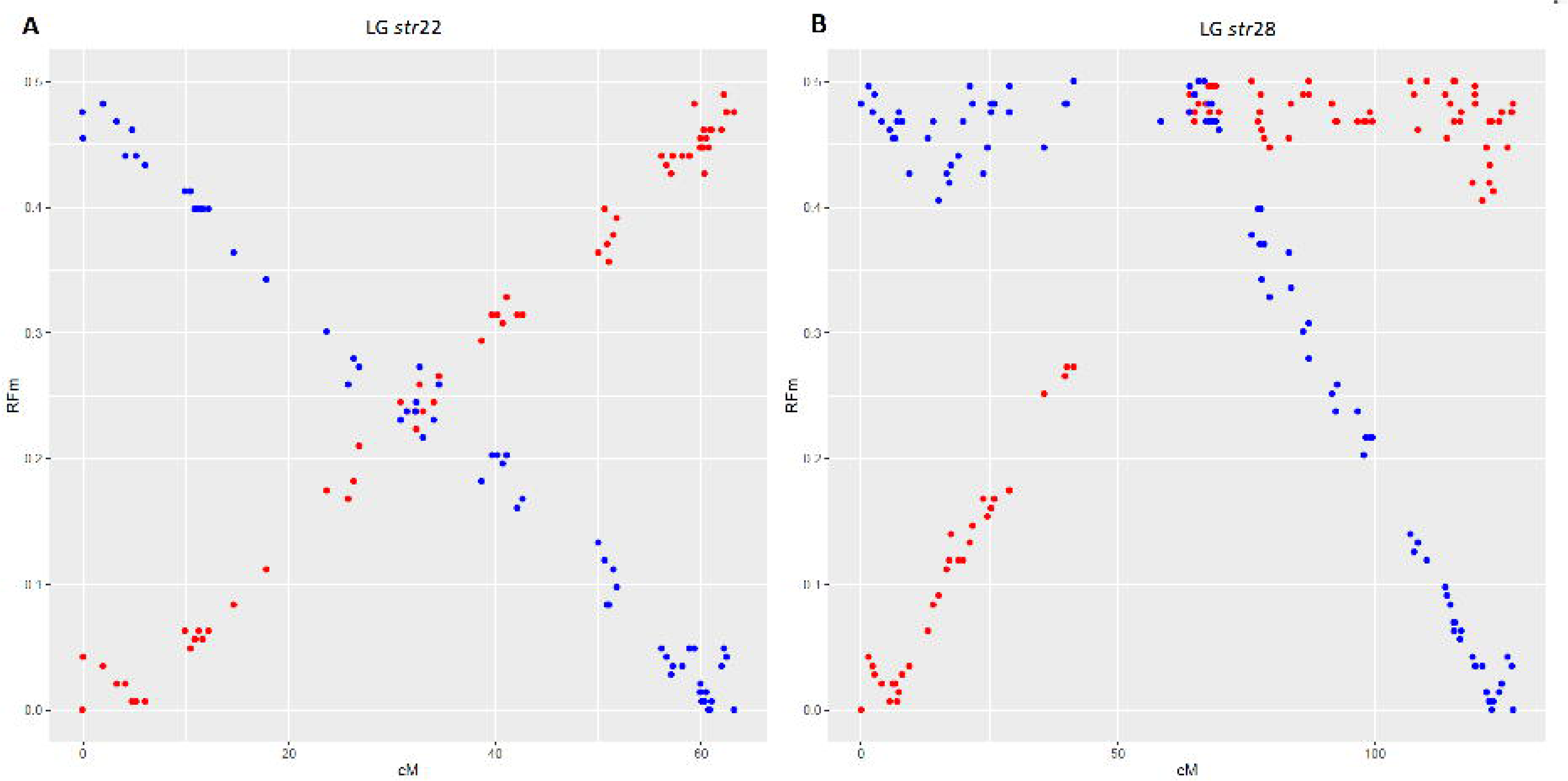
Plots illustrating the recombination frequency estimates (RFm) for intervals between markers along two different LGs. For each LG, RFm was calculated from both chromosomal extremities (right: red circles; left: blue circles), using each of the two terminal markers as a reference starting point. The RFm plot of (A) LG *str*22 illustrates a classical acrocentric pattern, while (B) LG *str*28 displays a classical metacentric pattern with a centromere position around 70 cM. The RFm plots of all brown trout LGs obtained in the two families are illustrated in Figure S1.

### Linkage map comparisons

A total of 87 microsatellites markers (82/160 from Gharbi *et al.* [2006], 5/5 from Leitwein *et al.* [2016] and the LDH locus [McMeel *et al.* 2001]) were successfully mapped to the Atlantic salmon genome, and anchored to the new brown trout linkage map using their pairing with RAD markers, following the method developed in Sutherland *et al.* (2016) (Figure 1; red dots). The correspondence between the microsatellite and the new RAD linkage map is provided in Table S2.

The identification of orthologous chromosomes arms between *S*. *trutta* (40LGs) and *S*. *salar* (29LGs) revealed frequent chromosomal rearrangements between the two species (Figure 3, Table 1). We found 15 one-to-one ortholog pairs (e.g., *Ssa*6-*Str*5, *Ssa*28-Str14; Figure 3, Table 1), 12 cases where salmon chromosomes correspond to either 2 or 3 LG in the brown trout (e.g., *Ssa*1, *Ssa*2; Figure 3), and only two cases where brown trout LGs correspond to two salmon chromosomes (*Str*1 and *Str*15; Figure 3). Local inversions between *S*. *salar* and *S*. *trutta* genomes were also observed as for instance *S*. *salar* LG6 and *S*. *trutta* LG5 (Figure S2). Apart from these inter-and intra-chromosomal rearrangements, a strongly conserved synteny was observed between the brown trout and the Atlantic salmon genomes.

**Fig. 3.**
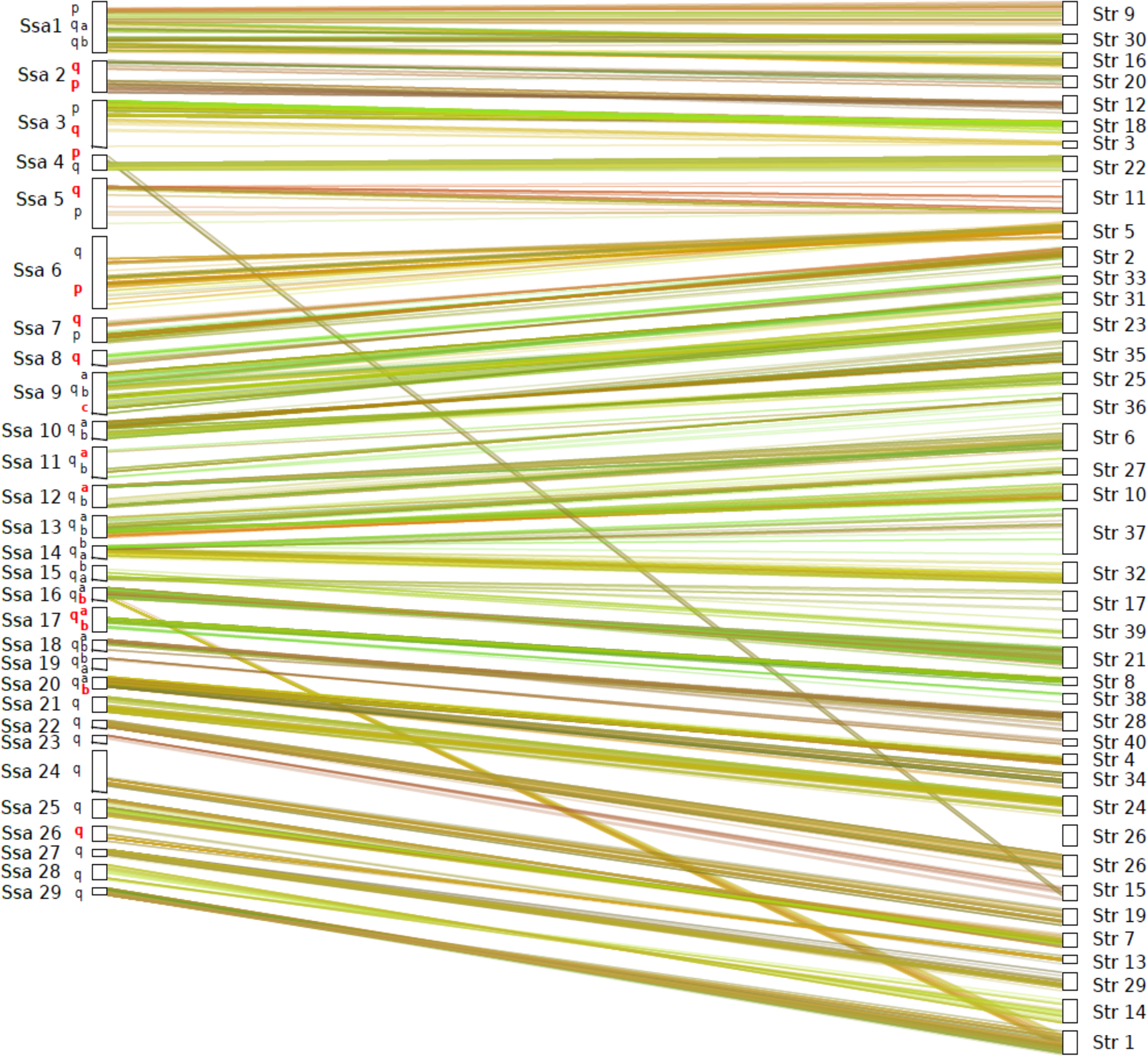
Dual synteny plot showing conserved syntenic blocks and chromosome rearrangements between *Salmo salar* (*Ssa*) and *Salmo trutta* (*Str*) LGs. Chromosomes arms (p, q_a_ and q_b_) are specified for *S*. *salar* LGs, and the seven pairs of chromosome arms that still exhibit residual tetrasomy in salmon appear in red (Lien et al., 2016).

Comparison with the brook charr linkage map revealed orthologous genomic regions with the brown trout and the Atlantic salmon (Table 1), thus allowing for the identification of ancestral and derived chromosomes structures in the *Salmo* genus. We found 8 chromosomal rearrangements that occurred in the ancestral lineage of the *Salmo* genus before speciation between *S*. *trutta* and *S*. *salar*, including 5 events of fusions (e.g., *Sfon*31-*Sfon*14, Table 1 and 2) and 3 events of fission (e.g., Sfon08; Table 1 and Figure 4). While no rearrangement was detected in the *S*. *trutta* branch, the Atlantic salmon branch contained 13 events of fusion and 2 events of fission (Figure 4). Among these fusion events, note that *Str*17 and *Str*39 are related to *Ssa*15, which was previously suspected to effectively be the result of a fusion (Phillips *et al.* 2009), but which is also known to host *SdY*, the male-determining system of the Atlantic salmon (Li *et al.* 2011) and most salmonids (Yano *et al.* 2013). As the microsatellite locus *OmyRT5TUF* was consistently shown to be linked to the *SdY* locus in the brown trout (Gharbi *et al.* 2006; Li *et al.* 2011), it may indicate that LG *Str*39 could support the male determining system of brown trout as the *OmyRT5TUF* locus was detected on this LG (Table S2). This has to be investigated further.

**Fig. 4:**
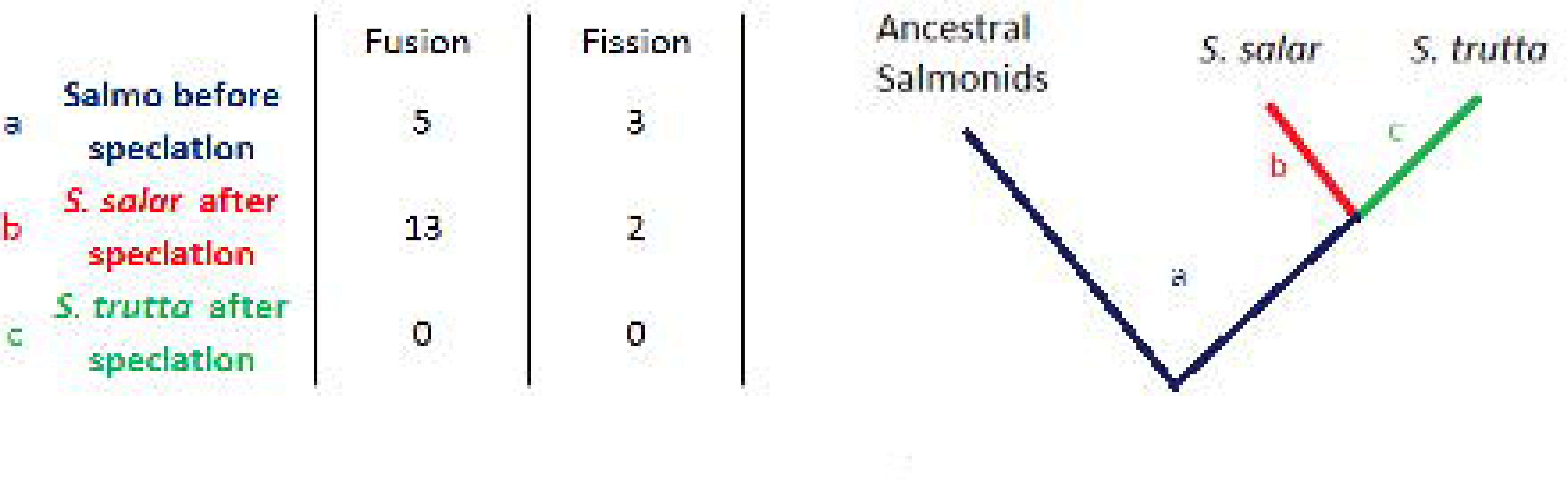
Summary of fission and fusion events within the *Salmo* genus, that were oriented using the brook charr (*Salvelinus fontinalis*) as an outgroup species. Rearrangements are classified with regards to their occurrence (a) before the *S*. *salari*/*S*. *trutta* speciation, (b) in the *S*. *salar* branch after speciation, or (c) in the *S*. *trutta* branch after speciation.

We estimated the average net nucleotide divergence between *S*. *trutta* and *S*. *salar* to *d*_a_ = 0.0187 (with π_b_= 0.0228, and π_w_= 0.0041).

### Estimations of genome-wide and local recombination rates

The brown trout genome-wide recombination rate averaged across the 3,721 markers anchored on the Atlantic salmon reference genome (RAD markers mapped to ungrouped scaffolds were discarded), was estimated at 0.88 ± 0.55 cM/Mb, and ranged from 0.21 cM/Mb (LG27) to 4.10 (LG 37) across linkage groups (Table 1, Table S1). Chromosomal variation in local recombination rate was observed for most chromosomes (e.g., LG *str*9 and LG *str*23, Figure 5). A significant positive correlation between local recombination rate and population nucleotide diversity was detected (R^2^ = 0.058; /*p*-value < 0.001; Figure 6).

**Table 1:**
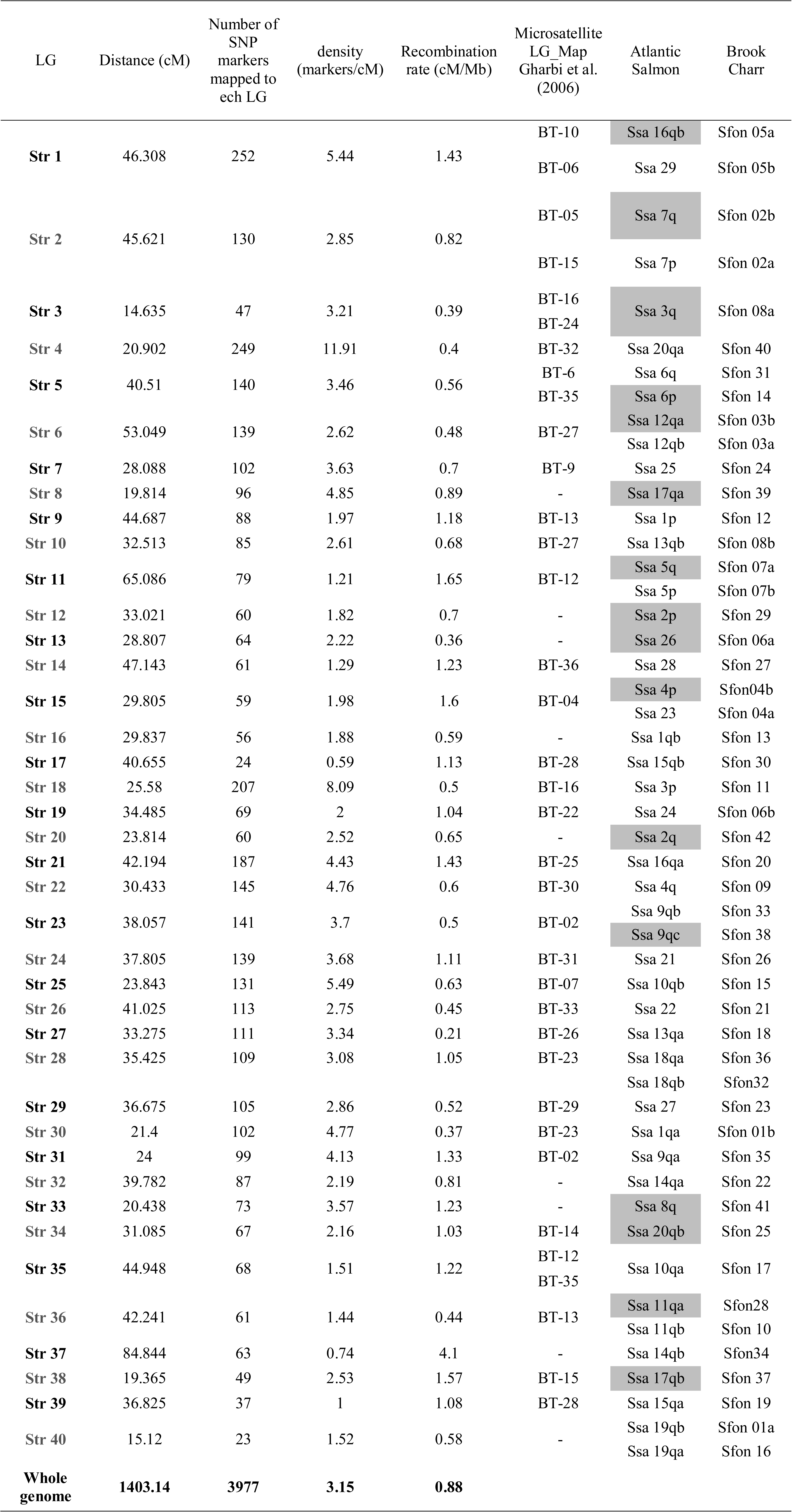
Summary of the main features of the brown trout high density linkage map based on the 3,977 mapped markers, including correspondence with the former generation linkage map established by Gharbi et al. (2006). The mean recombination rate is reported for each LG, as well as syntenic relationships with homologous chromosome arms from Atlantic salmon and brook charr. Grey-shaded arms correspond to the seven pairs involved in homeologous pairing (i.e. tetrasomic inheritance) in *S*. *salar* (Lien et al. 2016).

**Fig. 5.**
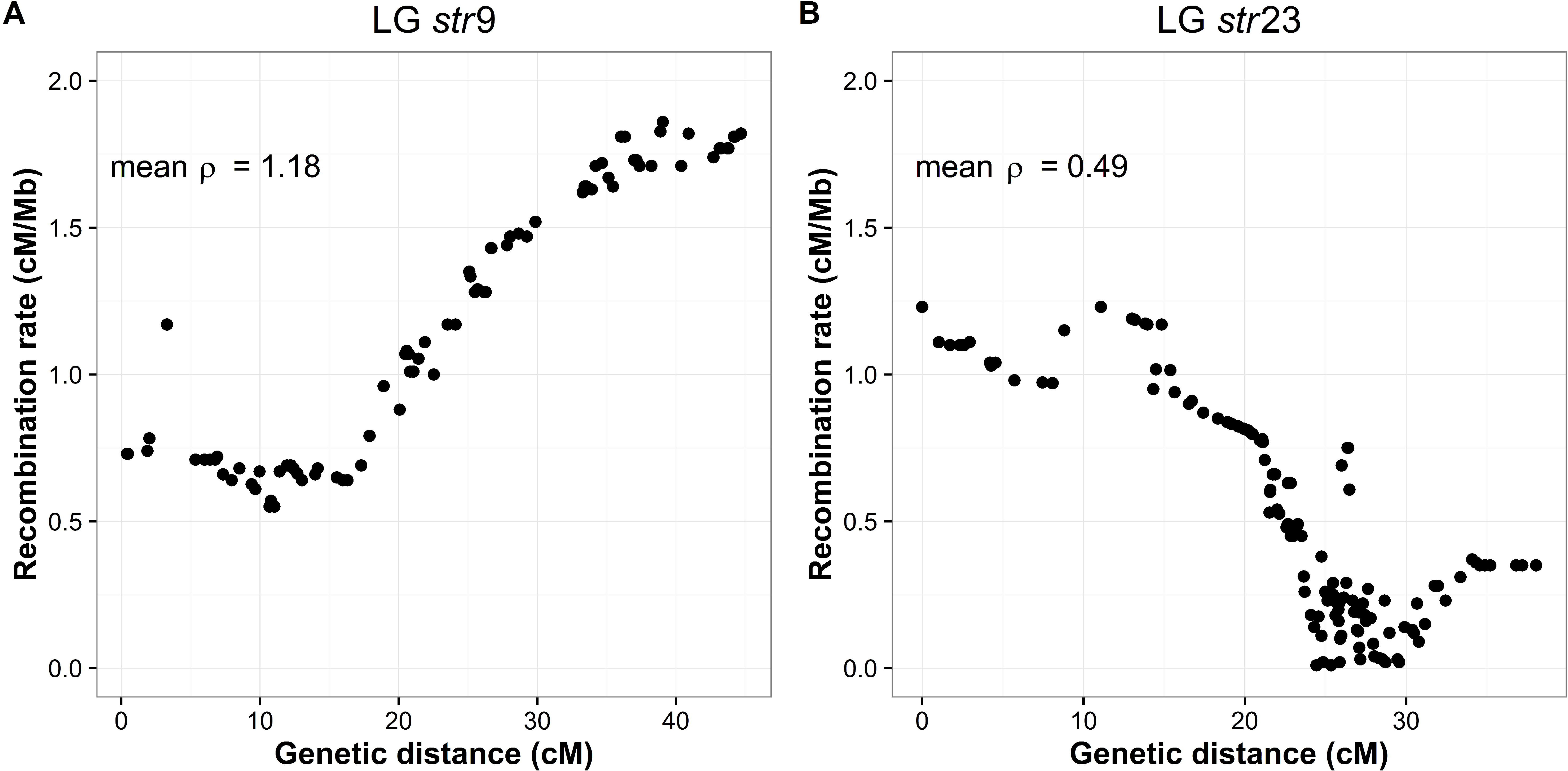
Estimates of local recombination rate (*ρ*) in cMIMb for markers located along two *Salmo trutta* linkage groups: (A) LG *Str*9 and (B) LG *Str*23. The average recombination rate (mean *ρ*) for each LG is given at the top left side of each figure. Local recombination rate estimates for other *S*. *trutta* LGs are reported in Table S1 and the average recombination rate of each LG is provided in Table 1.

**Fig. 6.**
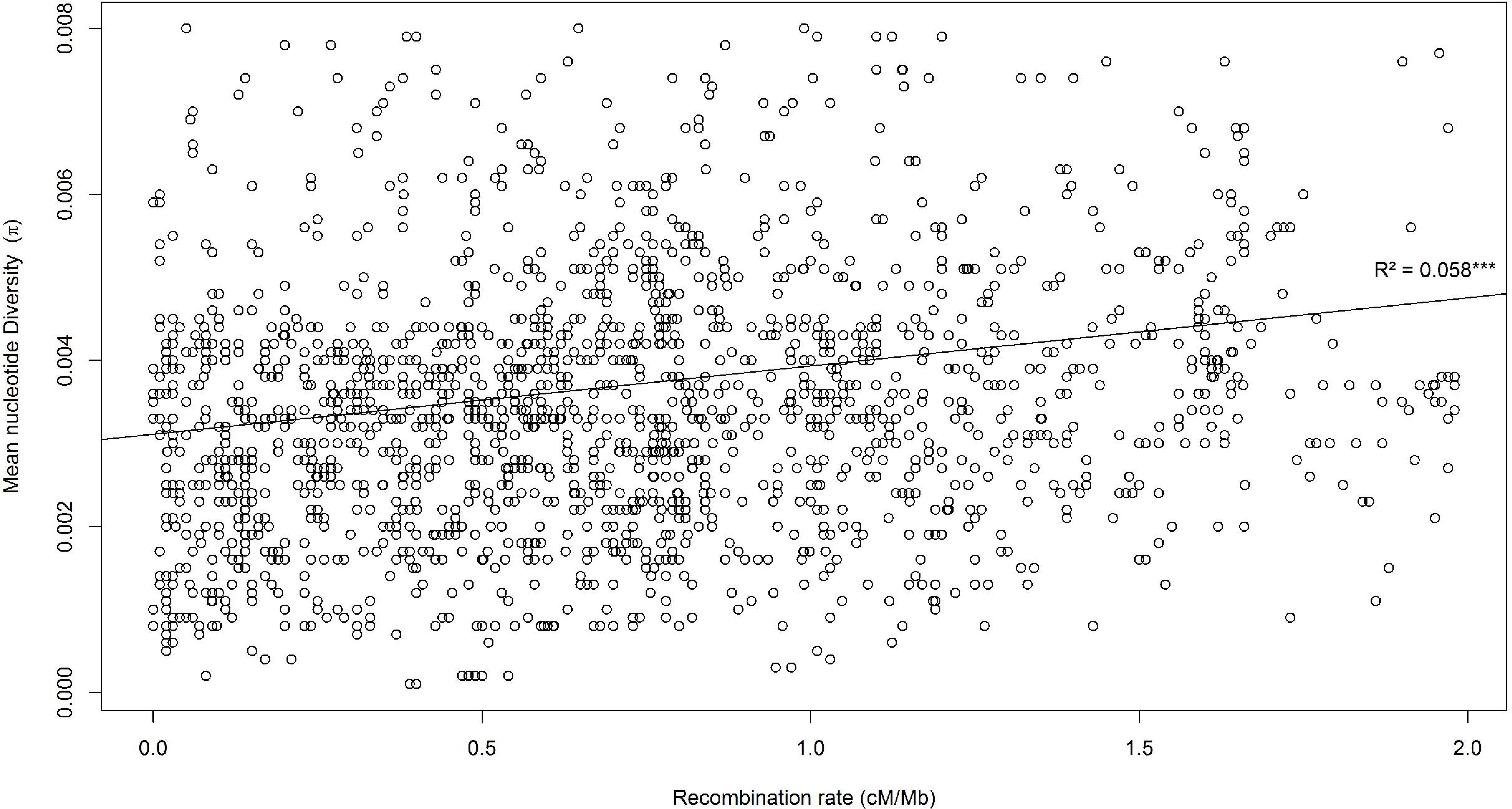
Genome-wide positive correlation between the mean nucleotide diversity in the Atlantic brown trout lineage (π) and the local recombination rate (*ρ*, in cMIMb), based on 3,038 SNP markers (R^2^ = 0.058; *p*value <0.001).

## DISCUSSION

This study describes the first high density linkage map in the brown trout *Salmo trutta*, which extends the microsatellite linkage map already available in this species (Gharbi *et al.* 2006). The map covered 1,403 cM and encompasses 3,977 non-distorted markers distributed across 40 linkage groups for which their centromere location was determined. Results showed consistent chromosome number and morphologies compared to the species’ karyotype (2*n*=80, NF=100; Phillips and Ráb 2001). High resolution mapping (3.15 markers per cM) was insured by the use of two mapping families each containing 150 offspring, and the number of markers informative in both families was close to the limit over which saturation is expected (i.e., 4,200 markers). The homogeneous distribution of makers across the genome, and the estimation of averaged recombination distances between both evolutionary lineages and sexes provide a highly valuable resource for future genomic, aquaculture, conservation and eco-evolutionary studies in brown trout. Beyond these general interests, this new linkage map also reveals strikingly different rates of chromosomal rearrangements that happened between the brown trout and the Atlantic salmon since their recent divergence.

### A new-generation RAD linkage map in brown trout

Comparison between the newly developed linkage map and the previous microsatellite map was made possible by the integration of 82 microsatellite markers from the former generation map (Gharbi *et al.* 2006). This enabled us to identify the correspondence between homologous LGs from the two maps, and to detect cases of LG merging (8/40) and LG splitting (9/40) in the microsatellite map compared to the RAD linkage map. Consistent with the reported overrepresentation of acrocentric compared to metacentric chromosome found in the brook charr (i.e., 8 metacentric and 34 acrocentric [Sutherland *et al.* 2016]), we found three metacentric morphologies against 32 acrocentric. This represents a total number of chromosome arms ranging from 86 to 96, which is lower but quite close to the 96-104 interval previously estimated in *S*. *trutta* (Martinez *et al.* 1991; Phillips and Ráb 2001). Our assessment of chromosome type did not perfectly match the results from Gharbi *et al.* (2006), since the three metacentric chromosomes identified here were found to be either acrocentric or undetermined in their study. However, the metacentric chromosomes *Str*2 and *Str*5 identified here had more than a single matching LG in the former microsatellite map, which suggests that these homologs represent the two telomeric regions of a larger metacentric linkage group which has been reconstructed in our study. On the contrary, the three metacentric chromosomes identified in Gharbi *et al.* (2006) appeared to be acrocentric using the phase information available from our dataset. This could be the consequence of pseudolinkage in the microsatellite map built using males of hybrid origin, in which the preferential pairing of conspecific homeologs followed by alternate disjunction may have generated an excess of non-parental gametes (Morrison 1970; Wright *et al.* 1983; Allendorf *et al.* 2015). In our study, the possible effect of pseudolinkage was avoided by using non-hybrid parents with a single lineage ancestry, thus possibly explaining the observed differences with the previous generation map.

Besides pseudolinkage, another broader consequence of past WGD in salmonids is the existence of duplicated loci that share the same alleles due to homeologous recombination in telomeric regions (Wright *et al.* 1983; Allendorf and Danzmann 1997; Allendorf *et al.* 2015). These residual tetrasomic regions challenge traditional linkage map reconstruction methods since duplicated loci (isoloci) tend to show Mendelian segregation distortion, and are therefore frequently excluded from linkage mapping analyses. Alternative approaches can be used to detect homeologous regions in experimental crosses (Lien *et al.* 2011; Brieuc *et al.* 2014; Waples *et al*. 2015). In brown trout, genome-wide patterns of RAD markers density and nucleotide diversity projected on the Atlantic salmon reference genome were also shown to provide indications about the location of duplicated regions (Leitwein et al 2016). Although our new RAD linkage map is based on non-distorted markers and therefore probably displays a reduced marker density in regions exhibiting tetrasomic inheritance, it does not completely exclude such regions. This is illustrated by the mapping of multiple RAD loci within the seven pairs of homeologs chromosome arms that retain residual tetrasomy in the Atlantic salmon genome (Lien *et al.* 2016) (Figure 3). Except for 3 of these 14 regions that received relatively few markers (*Ssa*3q, *Ssa*11qa, *Ssa*17qa on the salmon map; Figure 3), the remaining arms were well covered in our RAD linkage map, indicating that the filtering of non-Mendelian markers did not completely exclude the telomeric regions affected by homeologous crossovers. This feature of our RAD linkage map should insure a more even recombination rate estimation between sexes (Lien *et al.* 2011), since the commonly observed lower recombination rate in males compared to females in salmonids could be partly due to a poor characterization of telomeric regions in mapping studies (Allendorf *et al.* 2015). Thus, our final sex-averaged linkage map should not be affected by the integration of recombinational information from male meiosis, while providing at the same time more relevant estimates of recombination distances for future eco-evolutionary studies.

Following the same idea, integrating maps between parents of different lineage origins (Atlantic and Mediterranean) allowed us to generate a consensus linkage map in which the potential differences in recombination rates between these two allopatric lineages are smoothened. This is important for future conservation genomic studies, considering the extensive human-mediated admixture between Atlantic and Mediterranean brown trout populations in southern Europe (Berrebi *et al.* 2000; Sanz *et al.* 2006; Thaulow *et al.* 2014; Berrebi, 2015).

### The recent evolution of chromosomal rearrangements in the Salmo genus

Salmonids are still undergoing a rediploidization process since a WGD event that occurred around 60 Mya (Crête-Lafrenière *et al.* 2012; Near *et al.* 2012; Macqueen and Johnston 2014; Lien *et al.* 2016). This process, also described as functional diploidization by Ohno (1970), was accompanied by numerous chromosomal rearrangements within the salmonid family, making linkage maps of prime importance for studying the complex history of chromosomal evolution in salmonids (Sutherland *et al.* 2016). Several studies have pointed out a particularly high number of chromosome rearrangements in the lineage leading to the Atlantic salmon, involving mainly chromosome fusions that reduced the number of chromosomes to 29 in this species (Allendorf and Thorgaard 1984; Lien *et al.* 2016; Sutherland *et al.* 2016). However, the timing of these rearrangements remains largely unknown due to the lack of a closely related species in previous studies. Because *S*. *trutta* is the most closely related species to *S*. *salar* among salmonids, the new brown trout RAD linkage map allowed for inferring the phylogenetic positions of the chromosome arm rearrangements that occurred within the *Salmo* genus. The ancestral state of *Salmo* chromosome structure was further resolved using the brook charr as an outgroup, as well as other salmonid species when necessary (Sutherland *et al.* 2016). This enabled us to identify 5 fusion events (e.g., *Sfon*31-*Sfon*14, Table 1) and 3 fission events (e.g., *Sfon*08a and b, Table1) that occurred before the Atlantic salmon/brown trout speciation. Thus, most fissions and fusion events observed in the Atlantic salmon genome (Sutherland *et al.* 2016), specifically occurred after the divergence between the two Salmo species. The net nucleotide divergence between *S*. *salar* and *S*. *trutta* was estimated to 1.87%, which indicates a recent divergence time between these two sister species (i.e., less than 5 My). This estimate of nucleotide divergence should be compared to the previously reported estimate by Bernatchez *et al.* (1992) that was approx. 6%. It also probably challenges the >10My divergence time between *S*. *salar* and *S*. *trutta* reported in Crête-Lafrenière *et al.* (2012), and this point should be investigated further. Whatever the timing of divergence, all of the inter-chromosomal rearrangements that occurred after the salmon/trout speciation happened in *S*. *salar*. Indeed, we did not observe any fusion/fission event within the *S*. *trutta* branch (Figure 4), whereas 15 rearrangements mostly involving fusions (13 fusions and 2 fissions) specifically occurred in the Atlantic salmon branch. These Robertsonian translocations explain the increased number of metacentric chromosomes in the Atlantic salmon compared to the brown trout, which were produced by the recent fusion of acrocentric chromosomes (*Ssa*1 to *Ssa*4; Figure 3 and Table 1). However, fusion events could also result in the formation of acrocentric chromosomes in the Atlantic salmon (e.g., *Ssa*15; Figure 3 and Table1), thus explaining the apparent loss of chromosomes arms (Allendorf and Thorgaard 1984). Finally, intra-chromosomic rearrangements also occurred along the *S*. *salar* branch, such as, e.g., the inversion detected within *Ssa*6 and *Str*5 (Figure 3 and Figure S2). Altogether, these result support the previous views that many chromosomal rearrangements have occurred within the *Salmo* lineage (Allendorf and Thorgaard 1984; Sutherland *et al.* 2016), but demonstrate that most of them recently happened in the Atlantic salmon branch while at the same time *S*. *trutta* has retained a chromosomal architecture close to the ancestral salmonid form. The reasons for such a contrasted evolution of chromosome structure between these two species remain to be investigated. The fact that natural, viable hybrids occur between *Salmo* species (e.g., Jones, 1947; Garcia de Léaniz and Verspoor 1989) indicate that rearrangements have relatively poor consequences during meiosis, and in early life phases.

### Recombination rate variation across the brown trout genome

Despite extensive chromosomal rearrangements in the *S*. *salar* lineage, we could identify highly conserved synteny blocks within orthologous chromosome arms to reconstitute local alignments between the brown trout linkage map and the salmon physical map. One of our main objectives was to use these comparisons between maps to estimate local recombination rate variations along each linkage group in *S*. *trutta* (Figure 5). The average genome-wide recombination rate was estimated at 0.88 cM/Mb, however, with strong heterogeneity being found across the genome. High recombination rate regions were observed at extremities of some LGs (e.g., *Str*9; Figure 5), and genomic regions associated with very low recombination rates were also observed (e.g., *Str*23; Figure 5). In order to provide an indirect assessment of this local recombination rate estimation, for future eco-evolutionary studies, we verified that the frequently observed positive correlation between local recombination rate and nucleotide diversity (Corbett-Detig *et al.* 2015) also occurs in brown trout. In a comparative study across a wide range of species, Corbett-Detig *et al.* (2015) showed that the strength of this correlation is highly variable among species, being generally stronger in species with large *vs* small census population size. Although brown trout populations are usually considered to have relatively small census sizes (Jorde and Ryman 1996; Palm *et al.* 2003; Jensen *et al.* 2005; Serbezov *et al.* 2012), a significantly positive correlation was found between local recombination rate and nucleotide diversity using polymorphism data from the Atlantic lineage. Thus, recombination rate variation across the brown trout genome shapes the chromosomal landscape of neutral genetic diversity by modulating the efficiency of selection (both positive and negative) at linked neutral sites.

Variation in local recombination rate across genomic regions not only affects genome-wide variation patterns of polymorphism, but also has profound influence on a range of evolutionary mechanisms of prime importance for eco-evolutionary studies in brown trout. This recombination map should therefore have beneficial outcomes for our understanding of inbreeding depression, local adaptation, and the historical demography of natural populations (Rezvory *et al.* 2007; Dukić *et al.* 2016; Racimo *et al.* 2015). It will also be of precious help for understanding genome-wide introgression patterns in natural populations that have been restocked with individuals from a different lineage. Finally, a detailed picture of recombination rate variation in brown trout will help for choosing SNPs for the design of genotyping arrays in future association mapping experiments (e.g., Bourret *et al.* 2013; Houston *et al.* 2014).

### Conclusion

A high density sex- and lineage-averaged consensus linkage map for *S*. *trutta* was developed and the chromosome type (acrocentric or metacentric) of most linkage groups was identified. Interspecies comparisons with the Atlantic salmon allowed identifying genomic regions of highly conserved synteny within chromosome arms and contribute towards a better understanding of recent genome evolution following speciation in the genus *Salmo*. A recent and strong acceleration in the rate of chromosomal rearrangements was detected in the *S*.*salar* branch. These results should stimulate further research toward understanding the contrasted evolution of chromosome structure in sister species with less than 2% net nucleotide divergence. Recombination rate variation was found to influence genome-wide nucleotide diversity patterns in brown trout. These new genomic resources will provide important tools for future genome scans, QTL mapping and genome-wide association studies in brown trout. It also paves the way for a new generation of conservation genomics approaches that will look at the fitness consequences of introgression of foreign alleles into wild populations at the haplotype level.

## ACKNOWLEDGEMENTS

Authors wish to thanks E. Ravel and people from the Fédération Départementale de la Pêche de l’Hérault (France) and of the Babeau hatchery for their help at several steps of this project. We would like to thanks Y. Anselmetti for his kind advice on bioinformatics computing. Many thanks to M. Limborg for his helpful scripts and advice in particular for the centromere location. We thanks B. Sutherland for his suggestions with MapComp. This project largely benefited from the Montpellier Bioinformatics Biodiversity cluster computing platform. Presequencing steps necessary to the production of SNP data were performed at the Genotyping and Sequencing facility of the LabEx CeMEB (Centre Méditerranéen pour l’Environnement et la Biodiversité, Montpellier), and RAD sequencing performed at MGX (Montpellier GenomiX facility, Montpellier, France, http://www.mgx.cnrs.fr/). ML was partly supported by a grant of the LabEx CeMEB.

